# Heroin Choice Depends on Income Level and Economy Type

**DOI:** 10.1101/788703

**Authors:** Tommy Gunawan, Yosuke Hachiga, Christopher S. Tripoli, Alan Silberberg, David N. Kearns

## Abstract

**Rationale:** In a previous study investigating choice between heroin and a non-drug alternative in animals, reductions in income (i.e., choices/day) caused the percentage of income spent on heroin to progressively decrease. In contrast, another study found that humans with opioid use disorder spent the majority of their income on heroin even though they had little income. Comparison of these two studies suggests that the seemingly conflicting results could be explained by differences in the underlying economy types of the choice alternatives.

**Objective:** The present experiment tested the hypothesis that the effect of income changes on choice between heroin and a non-drug alternative depends on economy type.

**Methods:** Rats chose between heroin and saccharin under three income levels. For the Closed group, the choice session was the only opportunity to obtain these reinforcers. For the Heroin Open group and the Saccharin Open group, choice sessions were followed by 3-h periods of unlimited access to heroin or saccharin, respectively.

**Results:** As income decreased, the Closed and Heroin Open groups, but not the Saccharin Open group, spent an increasingly greater percentage of income on saccharin than on heroin. The Saccharin Open group, compared to the other groups, spent a greater percentage of income on heroin as income decreased.

**Conclusions:** Results confirm that the effects of income and economy type can interact and this may explain the apparently discrepant results of earlier studies. More generally, findings suggest that situations where heroin choice has little consequence for consumption of non-drug alternatives may promote heroin use.

---

Addiction has been conceptualized as persistent choice of drugs over non-drug alternatives (Ahmed et al. 2013; Banks and Negus, 2017; Bickel et al. 2011; Heyman 2013). Indeed, the diagnostic criteria for substance use disorder emphasize increased allocation of behavior towards drugs at the expense of other activities (Banks and Negus 2012). Identifying factors that promote choice of drugs over non-drug alternatives will help understand why persistent drug use occurs and may suggest treatment approaches that reduce drug use. According to behavioral economics, income is an important determinant of choice between two goods (Hursh 1980). If consumption of a good increases with increases in income, the good is defined as a normal good. These can be further characterized as necessity goods or luxury goods (Colander 2012). For necessities, reductions in income produce relatively small decreases in consumption; for luxuries, reductions in income produce relatively large decreases in consumption. Therefore, as income decreases, the proportion of income allocated to necessities increasingly rises (Goolsbee et al. 2016). The distinction between necessities and luxuries is not an absolute one; rather, there is a continuum on which goods fall in this regard (Lea 1978).

In a classic study, Elsmore et al. (1980) investigated the effect of income changes on choice between heroin and food in baboons. Income, operationalized as the number of reinforced choices subjects could make per day, was manipulated by varying the length of the inter-trial interval between choices. In the highest income condition, baboons could make 720 choices over the course of a 24-h session, while in the lowest income condition they could make 120 choices. Elsmore et al. found that reducing income produced large decreases in heroin consumption but had relatively little effect on food consumption. Consequently, the proportion of income spent on food increasingly rose while the proportion spent on heroin progressively fell as income was reduced. This pattern of results suggests that for baboons, heroin is a luxury good and food is a necessary good. Studies investigating the own-price elasticity of demand for heroin in rats have led to similar conclusions (Gunawan et al. 2019; Schwartz et al. 2017).

The finding by Elsmore et al. that baboons with low income spent only a small proportion of income on heroin seems inconsistent with what has been observed in humans with opioid use disorder. For example, Roddy and colleagues (Roddy and Greenwald 2009; Roddy et al. 2011) surveyed a group of heroin-dependent individuals and found that although they had very low income (about $723/month in wages), they spent about three quarters of their income on heroin. Subjects reported that family and friends were relied on for basic living expenses (e.g., food, shelter, etc.). When asked how their heroin use would be affected if family and friends no longer subsidized living expenses, subjects indicated that they would reduce heroin purchases by half (Roddy et al. 2011). The sensitivity of heroin consumption to the availability of subsidies provided by family and friends suggests a potential explanation for the seemingly conflicting results observed by Elsmore et al. and Roddy et al. Specifically, the economy for the non-drug alternative to heroin was closed in the Elsmore et al. study but the economy for non-drug alternatives was open for subjects in the Roddy et al. study.

For the baboons in the Elsmore et al. study, the only way to obtain food was by spending some of their income on it during the choice session. Total food intake depended completely on subjects’ choice behavior. The economy for food, therefore, met the definition of a closed economy (Hursh 1980). Likewise, the economy for heroin was closed. Subjects in the Roddy et al. study, on the other hand, could obtain food and shelter by spending their own income, but they could also rely on subsidies from family and friends for these expenses. Their consumption of these non-drug alternatives was therefore largely independent of their spending behavior. When there is a degree of independence between total consumption of a good and the subject’s behavior, the economy for that good is open (Hursh 1980). The difference in the economy type for the non-drug alternative(s) across studies may explain the difference observed in the allocation of income to heroin when income was low.

The foregoing analysis suggests a testable hypothesis: the effect of income on choice depends on the underlying economy types of the choice alternatives. When given a choice between two reinforcers and both are available in a closed economy, reductions in income should lead to an increase in the allocation of behavior towards the reinforcer that is considered a necessity. This is what Elsmore et al. found. However, if the economy for the necessity is open while the economy for the luxury is closed, income reductions may no longer produce the usual effect. Because the subject can obtain the necessity without having to spend income on it, income can be spent on the luxury without decreasing total consumption of the necessity. Therefore, reductions in income are less likely to shift allocation of income towards the necessity when the economy for the necessity is open as compared to when it is closed.

The hypothesis that the effect of changes in income depend on economy type was tested here in an experiment where rats chose between heroin and saccharin. Previous studies in rats have found that demand for saccharin is less price elastic than demand for heroin, which suggests that saccharin is more like a necessity than is heroin for rats (Gunawan et al. 2019; Schwartz et al. 2017). Income was manipulated by varying the length of the interval between trials during daily 3-h choice sessions. When the interval was short (e.g., 20 s), rats could make many choices per session. When the interval was long (e.g., 10 min), rats had few opportunities to choose. Economy type was manipulated by varying the availability of post-choice-session heroin or saccharin. For rats in the closed economies group, rats’ only chance to obtain heroin or saccharin was by choosing it during the choice session. For rats in the open heroin economy group, heroin infusions were made available on a fixed-ratio (FR) 1 schedule after each choice session for a 3-h period. Rats in the open saccharin economy group had FR-1 access to saccharin reinforcers during the 3-h post-choice-session period.

It was predicted that when the economies for both reinforcers were closed, reducing income would cause an increasingly greater percentage of income to be allocated to saccharin than to heroin. Opening the heroin economy was expected to enhance this effect because heroin forgone during the choice session could be easily replaced during the free-access period after the choice session. Of most interest were the results of the group for which the saccharin economy was open. It was predicted that, compared to the other two groups for which the saccharin economy was closed, this group would allocate a greater percentage of choices to heroin as income decreased. Because this group always had free access to saccharin after the choice session, they could continue to spend income on heroin without consequence for their total consumption of saccharin.

## Method

### Subjects

Thirty-six adult male Long-Evans rats (Envigo, Frederick, MD), weighing approximately 300 g at the beginning of experimental sessions, completed the experiment. Five other rats began the experiment but were excluded before any of the primary data could be collected due to catheter failure (*n* =1), health problems (*n* = 2), failure to acquire heroin self-administration (*n* = 1), or failure to habituate to the self-administration tether apparatus (*n* = 1). Rats were individually housed in plastic cages with wood-chip bedding and had unlimited access to rat chow and water in their home cages. The colony room where the rats were housed had a 12-h light:dark cycle with lights on at 08:00 h. Training sessions were conducted five days per week during the light phase of the light:dark cycle. Throughout the experiment, rats were treated in accordance with the Guide for the Care and Use of Laboratory Animals (National Academy of Sciences 2011) and all procedures were approved by American University’s Institutional Animal Care and Use Committee.

### Apparatus

Training took place in 20 Med Associates (St. Albans, VT) operant test chambers. Each chamber measured 30.5 × 24 × 29 cm and had aluminum front and rear walls with clear polycarbonate side walls. Three Med-Associates retractable levers were located on the front wall of the chamber. Saccharin reinforcers were provided by operation of a Med-Associates retractable sipper tube and bottle containing a 0.2% saccharin solution. The aperture through which the sipper tube inserted was located above the middle lever. A 100-mA cuelight was located above the left and right levers. A speaker was located in the center of the front wall near the ceiling. A 100-mA houselight was located at the rear of the chamber near the ceiling. Heroin (provided by the Drug Supply Program, National Institute on Drug Abuse, Bethesda, MD) in a saline solution at a concentration of 0.1 mg/ml was infused at a rate of 6.5 ml/min by 20-ml syringes driven by Med-Associates (St. Albans, VT) syringe pumps. Tygon tubing extended from the 20-ml syringes to a 22-gauge rodent single-channel fluid swivel (Instech Laboratories, Plymouth Meeting, PA) and tether apparatus (Plastics One, Roanoke, VA) that descended through the ceiling of the chamber. Heroin was delivered to the subject through tubing that passed through the metal spring of the tether apparatus.

### Procedure

#### Acquisition of lever pressing for saccharin

All rats were first trained to press the right lever for saccharin reinforcers on a FR-1 schedule during sessions lasting 2 h. The houselight was illuminated for the duration of the session. Each lever press resulted in insertion of the sipper tube for 20 s. The cue light above the right lever was illuminated during this period. Once rats regularly pressed the lever, the duration of sipper tube insertion (and cue-light illumination) was reduced to 10 s. Rats were trained on the FR-1 procedure with 10-s saccharin sipper tube insertions for a minimum of 10 sessions and until they earned at least 30 reinforcers for three consecutive sessions.

#### Surgery

After meeting the saccharin lever-press acquisition criterion, rats were surgically prepared with chronic indwelling jugular vein catheters, using procedures described in detail elsewhere (Thomsen and Caine 2005; Tunstall and Kearns 2014). In brief, approximately 3.5 cm of Silastic tubing was inserted into the right jugular vein. From this insertion site, an additional 12 cm of Silastic tubing passed under the skin to the midscapular region where it connected to the 22-gauge stainless steel tubing of a backmount catheter port (Plastics One, Roanoke, VA) that was implanted subcutaneously. The spring tether in the chamber was attached to the threaded plastic cylindrical shaft of the port that protruded through an opening in the skin. All surgery was conducted under ketamine (60 mg/kg) and xylazine (10 mg/kg) anesthesia. Rats were given 7–10 days to recover from surgery. Catheters were flushed daily with 0.1 ml of a saline solution containing 1.25 U/ml heparin and 0.08 mg/ml gentamicin.

#### Acquisition of lever pressing for heroin

After recovering from surgery, all rats were trained to press the left lever for heroin on an FR-1 schedule during sessions lasting 2 h. Each press resulted in infusion of 0.03 mg/kg heroin and illumination of the cue light above the left lever for 10 s. Rats were trained on this procedure for 10 sessions.

#### Choice

After acquiring the heroin lever-press response, rats were assigned to one of three groups (*n* = 12 for each group): the closed economies group (Closed), the open heroin economy group (Heroin Open), or the open saccharin economy group (Saccharin Open). Assignment was made with the goal of matching groups in terms of numbers of saccharin reinforcers and heroin infusions obtained during the last three sessions of acquisition.

All groups received a daily 3-h choice session that began with insertion of the left and right levers, illumination of the houselight, and presentation of white noise (72-74 dB) that remained on for the duration of the session. The white noise was introduced at this stage to help rats in the open economy groups learn that during choice sessions (as opposed to previous sessions) they would have delayed access to heroin or saccharin. A press on the left lever resulted in a heroin infusion, retraction of both levers, illumination of the cue light above the lever for 10 s, and initiation of an inter-trial interval (ITI). The length of the ITI varied over phases, as described below. A press on the right lever resulted in insertion of the saccharin sipper tube, retraction of both levers, illumination of the cue light above the right lever, and initiation of an ITI. If a rat made no press within 2 min, the trial was scored an omission, both levers retracted, and the next ITI began.

Income was manipulated by varying the length of time between choices. In the high income condition, the ITI was 20 s, allowing for a maximum of 540 choices per session. In the moderate income condition, the ITI was 180 s, allowing for a maximum of 60 choices per session. In the low income condition, the ITI was 600 s, allowing for a maximum of 18 choices per session. Rats experienced each of the different income levels in separate phases. The order of the phases was counterbalanced such that half of the rats in each group were exposed the three income conditions in ascending order (low, moderate, high) and the other half experienced them in descending order (high, moderate, low).

For the first income level that rats were exposed to, training lasted for a minimum of eight sessions and until a stability criterion was met whereby for three consecutive sessions the proportion of heroin choices (heroin choices ÷ sum of heroin and saccharin choices) did not differ from the rolling three-session mean by more than 0.2. For the subsequent two phases, rats were trained to the same stability criterion, but the minimum number of sessions was reduced from eight to five. A larger minimum was used for the first phase to ensure that rats in the open economy groups had sufficient time to learn that there were always reinforcers available after each choice session.

For rats in the Heroin Open group, each choice session was followed by a 3-h period during which the middle lever inserted and presses on it were followed by a heroin infusion and 10-s illumination of the left cue light. The left and right levers remained retracted during this period. The Saccharin Open group was treated similarly except that, instead of receiving extra access to heroin, presses on the middle lever were followed by insertion of the saccharin sipper tube and illumination of the right cue light for 10 s. The Closed group also remained in the chamber for an additional 3-h period following each choice session, but all levers remained retracted and no reinforcers were available. White noise and the houselight remained on for all groups during the 3-h post-choice period.

#### Data analysis

The primary measure of interest was the percentage of income spent on heroin or saccharin. This was calculated by dividing the number of heroin or saccharin reinforcers by the maximum number of choices possible under a particular income condition. For example, if a rat made 15 heroin choices in the moderate income condition, which permitted a maximum of 60 choices, the percentage of income spent on heroin was 25%. Omissions and the absolute numbers of heroin infusions and saccharin reinforcers obtained during the choice session, and during the 3-h post-choice-session period for the open economy groups, were analyzed. Number of sessions to meet the lever-press acquisition criterion, as well as numbers of saccharin reinforcers and heroin infusions obtained during acquisition sessions, were compared across groups.

For all statistical tests, α was set to 0.05. Repeated measures, one-way, or mixed ANOVAs were used to test the significance of findings. Group was a between-subjects factor in ANOVAs. Income (high, moderate, low) and Reinforcer (heroin vs. saccharin) were within-subjects factors. In instances where a 3 × 3 (Group × Income) ANOVA indicated there was a significant interaction, separate 2 × 3 ANOVAs were performed for each of the three possible pairwise group comparisons to identify the source of the interaction. Paired-sample *t*-tests or Tukey posthoc tests were used, where appropriate, following significant *F*-tests. The Benjamini-Hochberg (1995) procedure was used to keep the Type 1 error rate ≤ 0.05 for collections of multiple related *t*-tests.

## Results

Rats in the Closed, Heroin Open, and Saccharin Open groups required means of 17.3 (± 1.0 SEM), 16.9 (± 0.9), and 19.0 (± 2.0) sessions, respectively, to meet the saccharin lever-press acquisition criterion (no group difference, *F* < 1). Averaged over the last three saccharin acquisition sessions, these groups obtained 78.1 (± 8.3), 76.9 (± 6.3), and 75.1 (± 7.2) reinforcers per session, respectively (no group difference, *F* < 1). The Closed, Heroin Open, and Saccharin Open groups then self-administered means of 15.1 (± 2.4), 15.8 (± 3.3), and 15.2 (± 2.2) heroin infusions, respectively, averaged over the last three heroin lever-press acquisition sessions (no group difference, *F* < 1). On the choice procedure, there were only six rats that needed more than the minimum eight or five sessions to meet criterion at each income level. No more than three additional sessions were ever required.

The left panel of Figure 1 shows the percentage of income spent on heroin at each income level for the three groups. In the high income condition, all groups spent approximately 3-4% of income on heroin. As income was reduced, however, group differences emerged. The Saccharin Open group increased the percentage of income spent on heroin to 42% in the low income condition. In contrast, the Closed and Heroin Open groups spent only 12% of income on heroin in the low income condition. The right panel of Figure 1 shows the percentage of income spent on saccharin. As income was reduced, the Closed and Heroin Open groups spent increasingly more of their income on saccharin than they did on heroin. In contrast, for the Saccharin Open group, the increase in the percentage of income spent on saccharin was similar to the increase observed for heroin. Across income levels, the Saccharin Open group spent a smaller percentage of income on saccharin than the other two groups.

**Figure 1.**
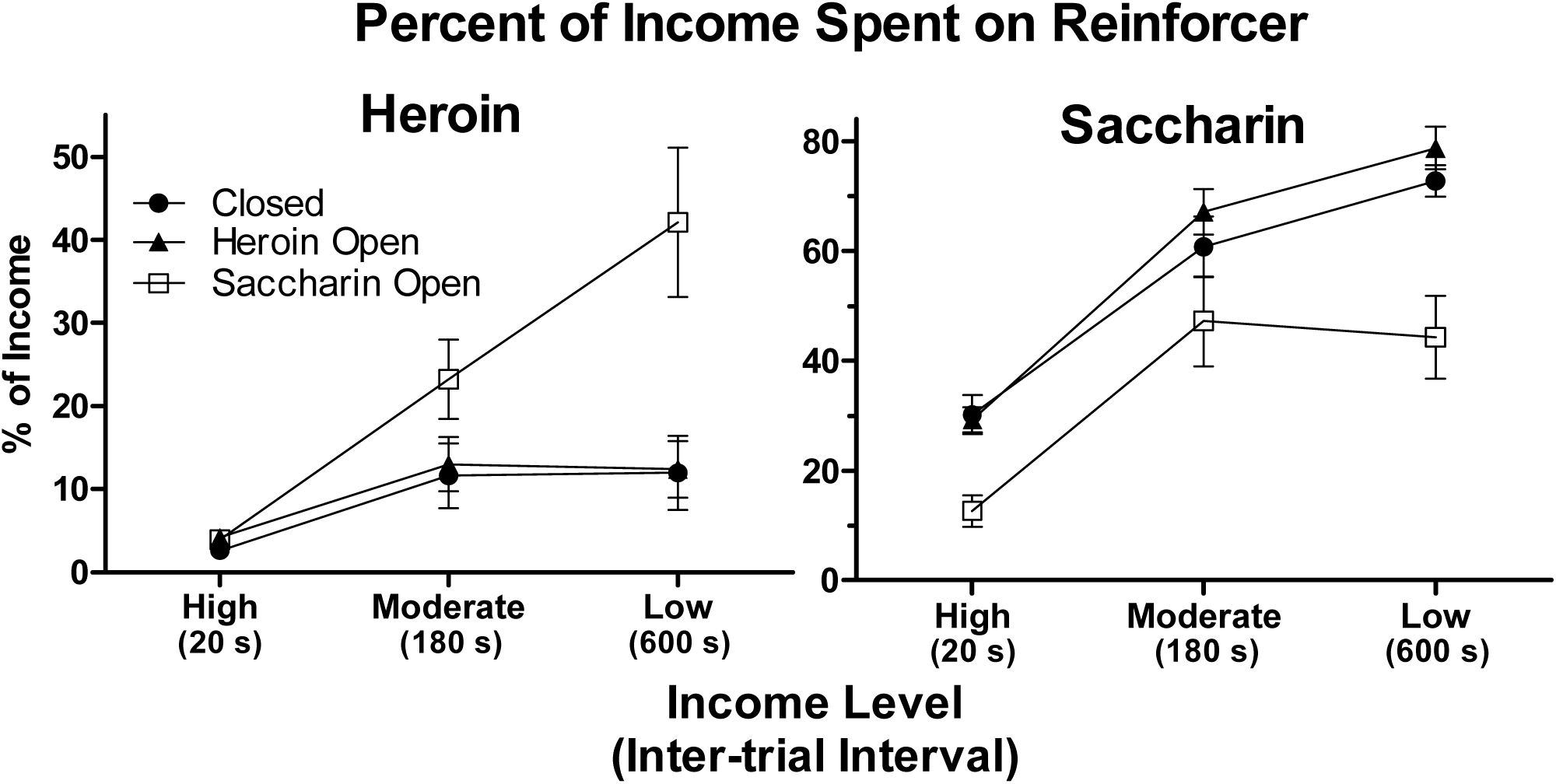
Mean (± SEM) percentage of income spent on heroin (left panel) and saccharin (right panel) as a function of income level (inter-trial interval) for the Closed (filled circles), Heroin Open (triangles), and Saccharin Open (squares) groups.

Statistical analyses confirmed these impressions. A 3 × 3 (Group x Income) ANOVA performed on the heroin data indicated there was a significant interaction (*F*[4,66] = 6.8, p < 0.001) as well as significant main effects of Group (*F*[2,33] = 5.9, *p* < 0.01) and Income (*F*[2,66] = 24.6, *p* < 0.001). Separate 2 × 3 (Group × Income) ANOVAs revealed that the interaction and effect of Group remained significant only when the Saccharin Open group was compared to the other two groups (interaction: both *F*[2,44]s ≥ 7.4, both *p*s < 0.005; Group: both *F*[1,22]s ≥ 7.0, both *p*s < 0.05), but not when the Closed and Heroin Open groups were compared to each other (both *F*s < 1). A 3 x 3 (Group x Income) ANOVA performed on the saccharin data from all three groups revealed significant main effects of Group (*F*[2,33] = 9.7, *p* < 0.001) and Income (*F*[2,66] = 123.5, *p* < 0.001), but no interaction (*F*[4,66] = 2.3, *p* > 0.05). Subsequent Tukey tests confirmed that, across income levels, the Saccharin Open group spent a smaller percentage of income on saccharin than the other two groups (both *p*s < 0.005). Additional 3 × 2 (Income × Reinforcer) ANOVAs performed on the data from each group separately indicated that there were significant interactions and main effects of Reinforcer for the Closed and Heroin Open groups (interaction: both *F*[2,22]s ≥ 14.3, both *p*s < 0.001; Reinforcer: both *F*[1,11]s ≥ 72.9, both *p*s < 0.001), but not for the Saccharin Open group (interaction: *F*[2,22] = 2.0, *p* > 0.15; Reinforcer: *F*[1,11] = 1.5, *p* > 0.2).

The top panel of Figure 2 shows that the number of heroin infusions self-administered by the Closed group decreased from about 14 to 2 as income was reduced from high to low. Compared to the Closed group, the Saccharin Open group consistently self-administered more heroin infusions, reducing from 21 infusions at the high income level to about 8 infusions when income was low. The Heroin Open group self-administered about the same number of heroin infusions as the Saccharin Open group during the choice session when income was high, but the number of infusions was reduced to about the same level as the Closed group under the moderate and low income conditions. A 3 × 3 (Group x Income) ANOVA performed on choice session infusions confirmed that there was a significant interaction (*F*[4,66] = 3.0, *p* < 0.05) as well as a significant main effect of Income (*F*[2,66] = 75.9, *p* < 0.001), but there was no main effect of Group (*F*[2,33] = 2.8, *p* > 0.05). A 2 × 3 (Group × Income) ANOVA performed on the Closed and Saccharin Open groups’ data indicated there were significant main effects of Group (*F*[1,22] = 4.5, *p* < 0.05) and Income (*F*[2,44] = 33.7, *p* < 0.001), but no significant interaction (*F* < 1). For the comparisons of the Heroin Open group with each of the other two groups, there was a significant Group x Income interaction (both *F*[2,44]s ≥ 3.3, both *p*s < 0.05) and a significant main effect of Income (both *F*[2,44]s ≥ 53.2, both *p*s < 0.001), but no main effect of Group (*F*[1,22] ≤ 2.6, both *p*s > 0.1).

**Figure 2.**
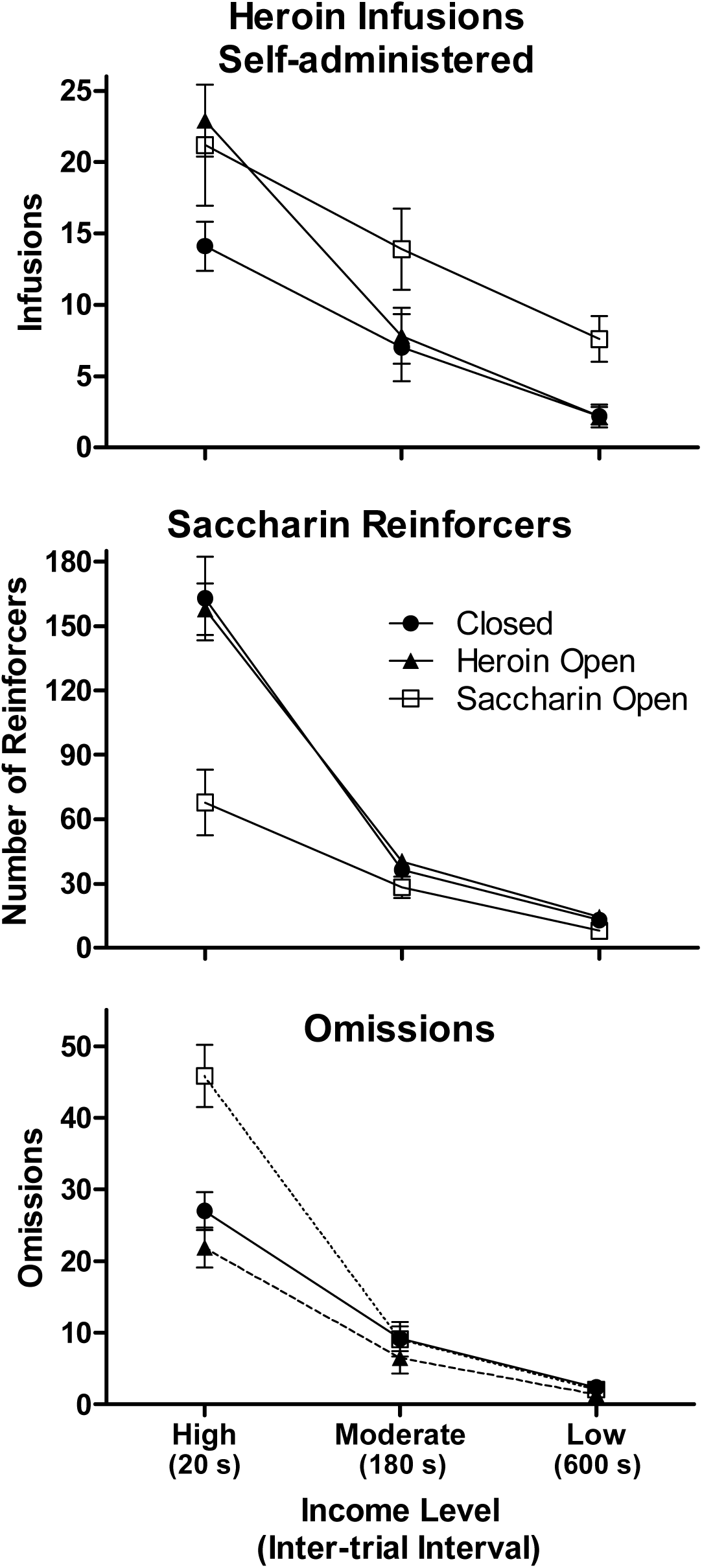
Mean (± SEM) numbers of heroin infusions (top panel), saccharin reinforcers (middle panel), and omissions (bottom panel) during choice sessions at each income level for the three groups.

The middle panel of Figure 2 shows the mean numbers of saccharin reinforcers obtained by the three groups during choice sessions at the three income levels. The Closed and Heroin Open groups both chose saccharin on about 160 trials during the high income condition, on about 40 trials during the moderate income condition, and on about 14 trials during the low income condition. In contrast, the Saccharin Open group only chose saccharin on approximately 70 trials during the choice session when income was high. This group performed similarly to the other two groups in the moderate and low income conditions. A 3 × 3 (Group × Income) ANOVA indicated that there was a significant interaction (*F*[4,66] = 10.7, *p* < 0.001) as well as significant main effects of Group (*F*[2,33] = 11.1, *p* < 0.001) and Income (*F*[2,66] = 154.7, *p* < 0.001). Separate 2 × 3 (Group × Income) ANOVAs revealed that the interaction and main effect of Group remained significant only when the Saccharin Open group was compared to the other two groups (interaction: both *F*[2,44]s ≥ 14.3, both *p*s < 0.001; Group: both *F*[1,22]s ≥ 13.6, both *p*s ≤ 0.001), but not when the Closed and Heroin Open groups were compared to each other (both *F*s < 1).

The bottom panel of Figure 2 shows that the Closed and Heroin Open groups were similar in terms of mean numbers of omissions, which decreased from approximately 20-25 in the high income condition to about 2 in the low income condition. The Saccharin Open group, in contrast, had about 45 omissions when income was high, but this number fell to approximately the same level as the other two groups under the moderate and low income conditions. A 3 × 3 (Group × Income) ANOVA confirmed that there was a significant interaction (*F*[4,66] = 14.3, *p* < 0.001) as well as significant main effects of Group (*F*[2,33] = 7.0, *p* < 0.005) and Income (*F*[2,66] = 215.1, *p* < 0.001). Subsequent 2 × 3 (Group x Income) ANOVAs indicated that the interaction and Group effect remained significant when the Saccarhin Open group was compared to the other two groups (interaction: both *F*[2,44]s ≥ 15.5, both *p*s < 0.001; Group: both *F*[1,22]s ≥ 5.7, both *p*s < 0.05), but not when the Closed group was compared to the Heroin Open group (interaction: *F* < 1; Group: *F*[1,22] = 2.0, *p* > 0.15).

The top panel of Figure 3 shows the numbers of extra heroin infusions or saccharin reinforcers obtained by the Heroin Open group and the Saccharin Open group, respectively, during the 3-h post-choice period. Both groups received about 50-65 reinforcers during this period across income levels. A 2 × 3 (Group × Income) ANOVA indicated that there were no significant effects of Income (*F* < 1), Group (*F* < 1), or their interaction (*F*[2,44] = 1.9, *p* > 0.15). The bottom panel of Figure 3 shows the numbers of heroin infusions or saccharin reinforcers obtained by these groups during the 3-h post-choice period over the final six blocks of three sessions regardless of income level. Thus, for half of the rats in each group, the final block was during the high income condition and for the other half it was during the low income condition. The final six blocks of sessions are shown because 18 was the minimum number of choice sessions that any rat had. Rats in the Heroin Open group escalated intake of heroin from approximately 35 infusions during the first block shown to about 55 infusions during the final block. In contrast, there was no escalation of saccharin intake in the Saccharin Open group. A 2 × 6 (Group × Block) ANOVA confirmed that there was a significant interaction (*F*[5,110] = 2.4, *p* < 0.05), but there were no main effects of Group (*F*[1,22] = 1.1, *p* > 0.25) or Block (*F* < 1). Subsequent repeated measures ANOVAs performed for each group separately revealed significant effects of Block for the Heroin Open group (*F*[5,55] = 3.2, *p* < 0.05), but not the Saccharin Open group (*F*[5,55] = 1.0, *p* > 0.4). Paired-sample *t*-tests confirmed that in the Heroin Open group, the numbers of infusions self-administered on Blocks 2-5 were significantly greater than the number self-administered on Block 1 (all *t*[11]s ≥ 2.3, all *p*s < 0.05).

**Figure 3.**
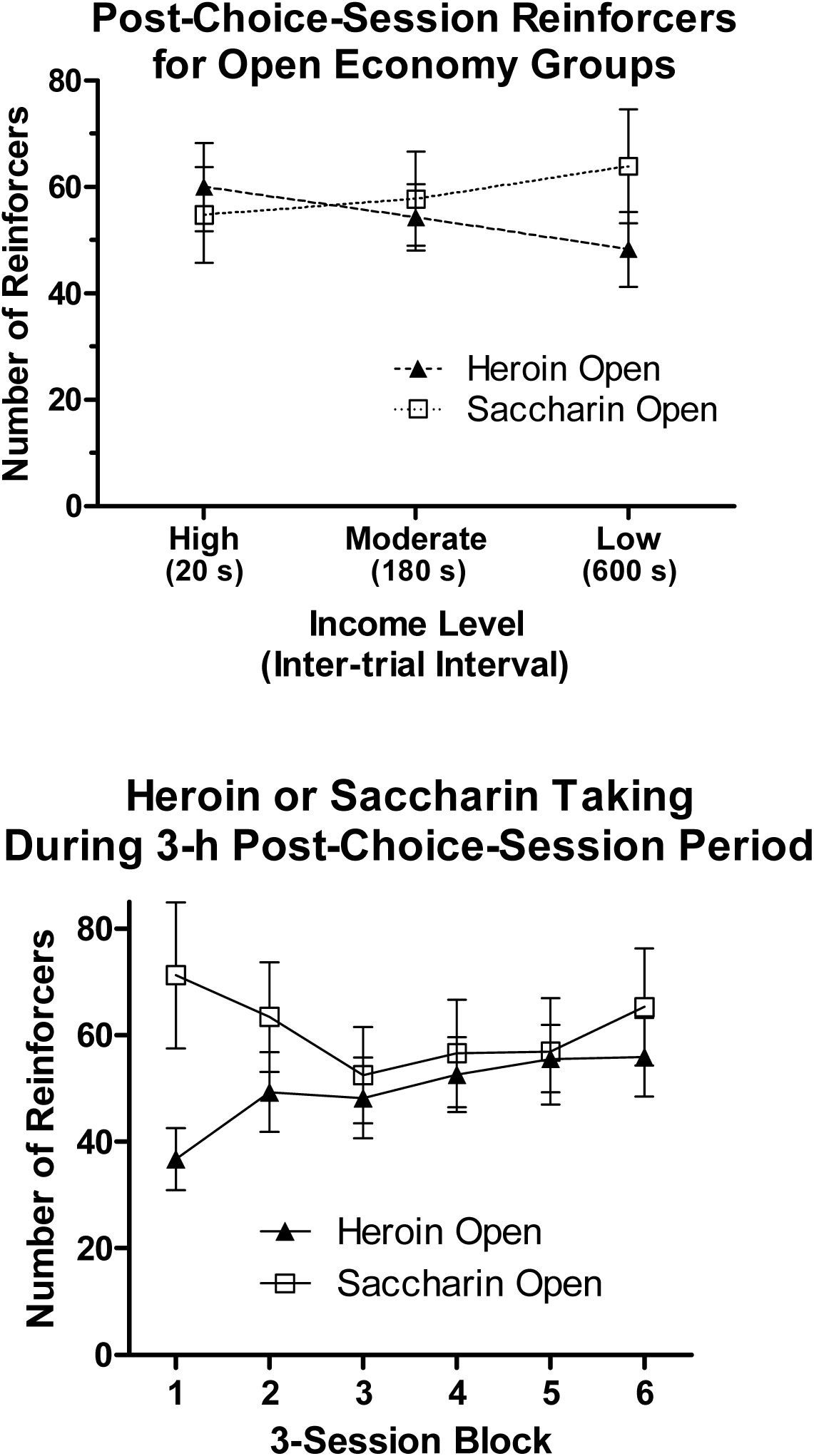
The top panel shows the mean (± SEM) numbers of heroin infusions or saccharin reinforcers obtained by the Heroin Open group and the Saccharin Open group, respectively, during the 3-h post-choice-session period at each income level. The bottom panel shows the mean (± SEM) numbers of heroin infusions or saccharin reinforcers obtained by these groups during the 3-h post-choice-session period over the last 18 sessions of training regardless of income level.

## Discussion

When the saccharin economy was closed, as it was for the Closed and Heroin Open groups, reductions in income caused an increasingly larger percentage of income to be spent on saccharin than on heroin. This shift towards saccharin in these groups is consistent with the effect of income reported by Elsmore et al. (1980) who studied choice between heroin and food in baboons in economies that were closed for both reinforcers. When the saccharin economy was open, as it was in the Saccharin Open group, three notable outcomes were observed. First, compared to the groups for which the saccharin economy was closed, the Saccharin Open group spent a greater percentage of income on heroin as income was reduced from high to low. Second, in contrast to the groups for which the saccharin economy was closed, reductions in income did not cause rats in the Saccharin Open group to spend an increasingly larger percentage of income on saccharin than on heroin. Third, the absolute numbers of heroin infusions self-administered by the Saccharin Open group was higher than that self-administered by the Closed group, especially when income was low. At the moderate and low income levels, there were 2- and 3-fold increases, respectively, in the numbers of heroin infusions self-administered by the Saccharin Open group compared to the Closed group.

In contrast to the effects of opening the saccharin economy, opening the heroin economy did not alter choice behavior. The Heroin Open group and the Closed group were remarkably similar with respect to how they allocated income to heroin and saccharin across income conditions. This lack of effect does not appear to be due to insufficient post-choice-session access to heroin. The Heroin Open and Saccharin Open groups did not differ in numbers of reinforcers obtained during the 3-h post-choice-session period. Further, the Heroin Open group escalated their heroin intake during this period over sessions, an effect that is typically only observed with a large amount of exposure to heroin (Vendrusculo et al. 2011). The lack of effect of opening the heroin economy here is the third instance where post-session access to self-administered intravenous drug failed to affect behavior during the session. Previously, we found that post-session access to cocaine (Kim et al. 2018) or to heroin (Gunawan et al. 2019) did not affect elasticity of demand for these drugs in experiments where, using the same procedures, post-session access to saccharin made demand for saccharin more elastic. These results suggest that, for rats, delayed drug reinforcers do not substitute for current drug reinforcers in the same way that delayed saccharin reinforcers substitute for current saccharin reinforcers. Such a result might be expected if, for example, delayed heroin is discounted at a higher rate than delayed saccharin. It is currently unknown whether this is the case in rats, but there is evidence that humans delay discount heroin more steeply than they do money (Madden et al. 1997, 1999).

It may be thought that the effects of opening the saccharin economy were due to between-session satiation. That is, rats in the Saccharin Open group might have drunk so much saccharin during the 3-h post-choice period that they were partly saccharin-sated at the start of the choice session the next day and this could have influenced the way they distributed their choices. There are reasons to doubt this satiation account. The half-life of saccharin in rats is only about 30 minutes (Renwick 1985; Sweatman and Renwick 1980), but there were at least 18 hours (more on weekends) from the end of the post-choice period on one day to the start of the next choice session. Furthermore, studies investigating factors controlling the termination of saccharin drinking in rats indicate that it is due to adaptation to the immediate orosensory stimulation provided by saccharin, rather than to delayed post-ingestive consequences, including fullness of the stomach with fluid (Mook et al. 1980, 1981). Finally, if saccharin satiation in the Saccharin Open group were driving results here, the largest group differences might be expected in the high income condition, where the most total saccharin consumption occurred. Instead, the Saccharin Open group diverged most from the other two groups in terms of heroin choice in the low income condition, when access to saccharin during the choice session was most restricted.

The rate of omissions was fairly high in the high income condition, especially in the Saccharin Open group. Instead of choosing more heroin or saccharin, rats often chose to not respond at all. Within-session satiation during choice sessions might explain the high rate of omissions in the high income condition. But in the moderate income condition, omissions still accounted for 10-15% of the total number of choices possible even though satiation was not likely since rats in all groups took less than half the numbers of saccharin reinforcers that they took in the high income condition. They could have more closely approximated the level of consumption of heroin and saccharin observed in the high income condition by making fewer omissions in the moderate income condition. Elsmore et al. (1980) also found a high rate of omissions in their baboons. In their high income condition, where up to 720 choices per day were possible, baboons only made a choice on 22% of trials. At the lowest income level, where 120 choices per day were possible, baboons only made a choice on 63% of trials. As Elsmore et al. noted, baboons in the low income condition could have maintained the level of heroin intake observed under the high income condition simply by making fewer omissions and without giving up any food. That they did not suggests that another behavior (e.g., sleeping, grooming, etc.) often successfully competed with food- or heroin-taking behavior. That was likely the case in the present experiment as well, at least during the high and moderate income conditions. Rats in all groups here rarely made omissions when income was low.

The results of the present experiment support the hypothesis that the effect of income on choice can vary depending on the underlying economy types of the choice alternatives. As a model of human behavior, the present results help to make sense of the finding in Roddy et al. (2011) that very low-income heroin users spent a large percentage of income on heroin. As the heroin users themselves reported, removing outside sources of non-drug alternatives would cause them to spend less of their income on heroin. More generally, these results suggest that situations where drug taking has little consequence for total consumption of non-drug reinforcement will promote drug taking. Conversely, situations that make total consumption of non-drug reinforcement strongly dependent on not using drugs will promote abstinence. Indeed, the success of contingency management treatment (for recent reviews, see Davis et al. 2016; De Crescenzo et al. 2018), which makes the contingency between drug use and loss of non-drug reinforcement explicit and consistent, accords with the view that drug taking, even in individuals with substance use disorder, is a behavior that is strongly determined by consequences. A full understanding of the relationship between drug taking and consequences will have to take economy type into account.

